# Domesticated pennycress is a predominantly self-pollinated crop

**DOI:** 10.64898/2026.04.08.716402

**Authors:** Tito Lavaire, Daniel McLaughlin, Shengjun Liu, Ray Kennedy, Thomas Sauer, Ratan Chopra, Kevin Cook

**Author notes:** Correspondence: Shengjun Liu; Kevin Cook.

## Abstract

Domesticated pennycress (*Thlaspi arvense* L.) is an emerging winter annual oilseed crop developed from field pennycress to deliver commercial agronomic performance, low-carbon feedstock for biofuel production, and value-added animal feed. Effective breeding, seed production, and stewardship of this newly domesticated crop requires a clear understanding of its potential for pollen-mediated outcrossing. Although prior evidence suggests that field pennycress is predominantly self-pollinated, direct experimental evidence remains limited. Here, we integrate laboratory, greenhouse, and field experiments to rigorously evaluate pollen viability, pollen movement, and outcrossing potential in domesticated pennycress. Pollen viability declined rapidly with increasing temperature and environmental exposure: pollen stored at 4°C reached 50% viability after 12 h, whereas pollen held at 37°C reached the same threshold after 2.5 h. Exposure to outdoor conditions with natural light and wind reduced viability to 50% within 1.6 h. Under greenhouse conditions, no cross-pollination was detected among plants flowering in close proximity within an agitated pollen-containment covering. Detectable cross-pollination occurred primarily when emasculated flowers were directly exposed to donor pollen, demonstrating that intact flowers strongly favor self-fertilization. Field trials using a trackable knockout gene edit provided no detectable evidence of pollen movement under field conditions across two locations, even at short distances from donor plots. Together, these results provide strong, multi-environment evidence that domesticated pennycress is predominantly self-pollinated and that outcrossing is rare under practical greenhouse and field production conditions.

## 1. Introduction

Domesticated pennycress (*Thlaspi arvense* L.) is a new winter annual oilseed crop developed from field pennycress, a commonly occurring non-crop species found in almost all areas of North America and most other continents (Phippen and Phippen, 2012; Chopra et al., 2018; Phippen, et. al., 2022; Basnet and Ellison, 2024). This new crop delivers the soil health benefits typical of a low-input cover crop (Dierberger et al., 2025), while also producing a low-carbon-intensity feedstock for renewable diesel and sustainable aviation fuel.

Traditional plant breeding in combination with advanced breeding techniques, such as marker-assisted breeding and gene editing, served to improve agronomic performance and seed compositional profiles in domesticated pennycress. Certain varieties of domesticated pennycress, sold as CoverCress®, have reduced erucic acid content, reduced fiber content, and reduced glucosinolate levels compared to field pennycress (Phippen et al.,2022; Tuite et al., 2025) that resulted from specific gene edits. These changes are analogous to the improvements in oilseed rape that resulted in canola (*Brassica napus* var. *napus* L.). Erucic acid can have adverse health effects on livestock when consumed at high levels. The lower fiber seed coat results in a thinner, yellow seed coat which is visually distinct from black to dark brown field pennycress grain with a lower acid detergent fiber content. The thinner, yellow seed coat also enables faster, more consistent germination and reduces the potential for dormancy compared with native field pennycress, thus reducing the potential for weediness (Mcginn et al., 2019). Reduced glucosinolates make the meal from crushed grain more palatable to livestock (Hartnell et al., 2023). During development, breeders selected and bred varieties exhibiting strong germination, timely flowering, lodging resistance, and timely harvest (Basnet et al., 2024; Azhar et al., 2025).

Understanding the extent to which domesticated pennycress may cross-pollinate with field pennycress is important for establishing guidelines to prevent outcrossing during breeding and seed production, in part to ensure the seed used for production will produce the expected grain product. Understanding the potential for outcrossing is unique to the plant in question (Stevens et al., 2004, Ingram, 2000; Maity et al., 2022), the United States Department of Agriculture (USDA) promulgated regulations implementing requirements of the Federal Seed Act that specify isolation requirements for common crop species grown from seed. Certified seed producers of listed crop species must comply with restrictions on what has been grown in a particular field in previous years and the distance from other sources of pollen in order to state that their seed is foundation, registered, or certified. Self-pollinated crops require less isolation than open pollinated crops (eCFR, 2026).

There are few published studies on pollen flow dynamics and outcrossing levels in pennycress. The available data support the conclusion that pennycress is primarily self-pollinated (Mulligan and Kevan, 1973, Sedbrook et al. 2014, Dorn et al., 2015). The floral structure, consisting of six stamens with four possessing anthers at the same height or slightly higher than the stigma, makes pennycress readily autogamous as the stamens curl inward as flowers mature (Warwick et al., 2002; Best and McIntyre, 1975). Sequence-based analysis by Dorn et al. (2015) revealed high levels of homozygosity within populations as evidence of self-pollination. Groeneveld and Klein, 2014, however, asserted that outcrossing is predominant and there are reports of insect pollinators (bees from the genus *Apis*, and the fly genera *Anthomyia* and *Pollenia*) actively foraging pennycress flowers (Al-Shehbaz, 1986). These disparities create uncertainty when developing protocols to prevent outcrossing under field conditions.

Groeneveld and Klein (2014) may have underestimated the rate of self-pollination as a result of the protocol they employed. The authors’ based their conclusion upon the observation that seed set of open-pollination and wind-pollination treatments was much higher than that of the self-pollination treatment, however their protocol design did not include a way to discern self-pollination from cross pollination on the same plant in the open and wind-pollination treatments. The self-pollination treatment involved covering flowers with an air-permeable pollen-exclusion bag, possibly leading to elevated temperatures near the flowers. The pollen exclusion bags relied upon 0.8mm perforations to enable air movement and would be impacted by greenhouse effects on warm days. Parker and Phippen (2012) reported an 84% reduction in pennycress pollen viability at 30°C and a 100% reduction at 32°C. Groeneveld and Klein (2014) did not report daily maximum temperatures from their experiments, nor did they measure the temperature inside the pollen exclusion bags. It is likely that temperatures within the bags exceeded levels sufficient to reduce pollen viability on warm, sunny days.

The experiments reported on herein further elucidate the fertilization process of pennycress flowers by characterizing the potential for outcrossing under both greenhouse and field conditions using a trackable marker system and reports on pennycress pollen longevity under different temperature and light conditions.

## 2. Materials and Methods

CoverCress Inc. developed all of the domesticated pennycress varieties used in these experiments.

### 2.1 Pollen viability kinetics

Pollen grains for each of the temperature and light treatments described were collected from three plants of the same genotype between 9 to 10 AM and placed on glass slides. A dissecting microscope allowed pollen grain visualization and removal, and harvested pollen was deposited on an uncovered glass slide. The fluorescein diacetate (FDA) procedure (Bou et al., 2009) differentiated viable and non-viable pollen grains. The FDA procedure relies on esterases in viable pollen to cleave acetate from FDA, resulting in fluorescein and providing a clear visual signal (Figure 1 provides an example). Briefly, a five-minute immersion in 250 μL FDA solution under dark conditions allowed dye to penetrate pollen on the slides and begin the reaction with pollen esterases. Twelve to 35 stained pollen grains were evaluated on each slide using a Leica DMI4000 B microscope (Leica CTR 4000 equipped with 100W halogen lamp for transmitted light and an external metal halide lamp to excite fluorophores, a DFC300FX camera and Leica application suite version 3.8 software). Images were collected using a Texas Red emission filter (BP 560/40). A sample of pollen heated on a glass slide for five minutes over a Bunsen burner served as a negative control. Manual counts of viable and non-viable pollen from 12 to 18 average views at each experimental time point were used to plot viability as a function of time and temperature after outlier removal.

**Figure 1.**
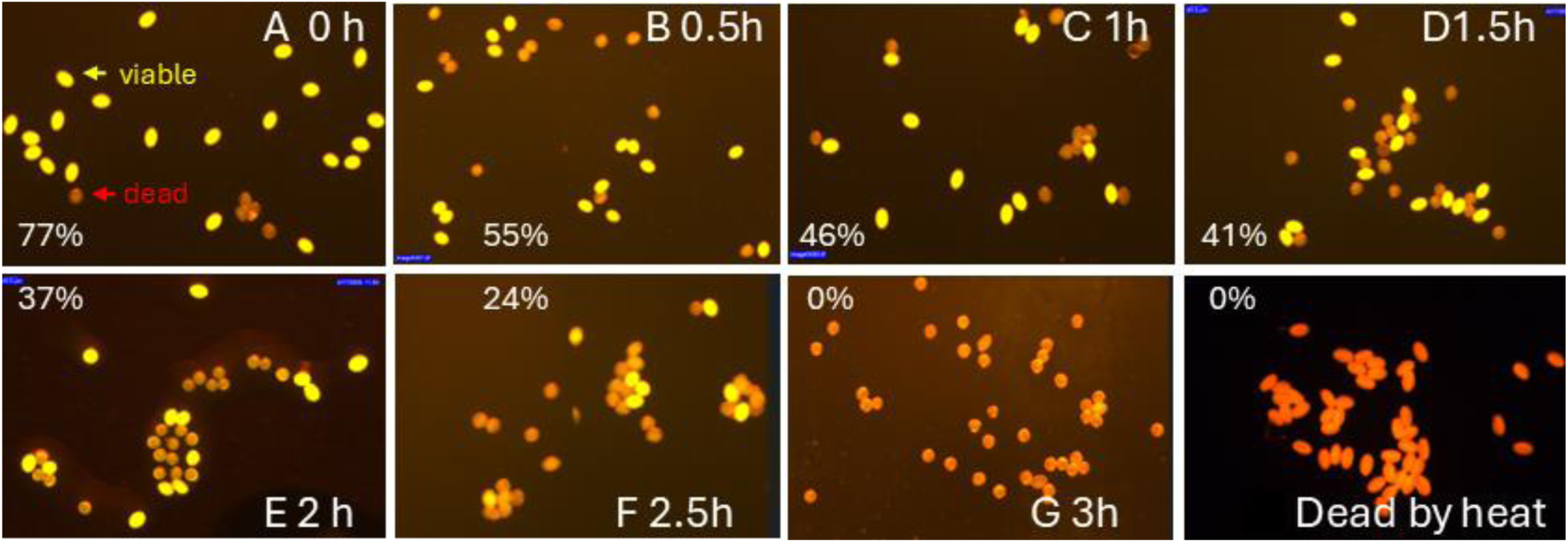
Viability of natural sunlight-treated domesticated pennycress pollen grains. FDA fluorescence of domesticated pennycress after 0 hours (Panel A, 20 viable pollen grains out of 26 evaluated =77% viable); 0.5h (B, 15/27=55%); 1h (C, 11/24=46%); 1.5h(D, 14/34=41%); 2h (E, 13/35=37%); 2.5h (F, 8/33=24%); 3h (G, 0/41=0%) under described outdoor conditions; Dead pollen standard after heat treatment(H).

For the experiments measuring viability in response to temperature, pollen samples were stored at 37 °C, 22 °C, and 4 °C for 0, 0.75, 1.5, 3, 6, 12, 24, and 36 hours after collection. To assess the impact of natural light and wind on viability, pollen samples on glass slides were placed on a screen table outdoors between the hours of 10:00am and 4:00pm (St. Louis, MO, on April 8, 2025, temperature range of 4.4-10 °C) with viability measurements 0, 0.5, 1.5, 2, 2.5, 3, 3.5, 4.5, 6, and 7 hours after collection.

### 2.2 Outcrossing Trials Under Greenhouse Conditions

#### 2.2.1 Close-proximity cross-pollination

Five transgenic families of the experimental pennycress line 183002-B-14 were created with Agrobacterium-mediated transformation via floral dip infiltration with the plasmid vector pARV532 containing an expression cassette for the DSRed2 (Red Fluorescent Protein, RFP) selectable marker, selecting putative single insertion plants, and self-pollinating the resulting hemizygous plants. These five T2 families segregated in a Mendelian fashion for expression of the DSRed2 cassette. Expression intensity enabled differentiation among RFP/RFP homozygous fixed (high intensity fraction), RFP/rfp hemizygous (moderate intensity fraction), and rfp/rfp null segregant plants (non-expressing). Figure 2 provides examples of the appearance of the three types under natural and detection lighting. Two homozygous fixed and two null seeds were selected from each family to evaluate pollen movement.

**Figure 2.**
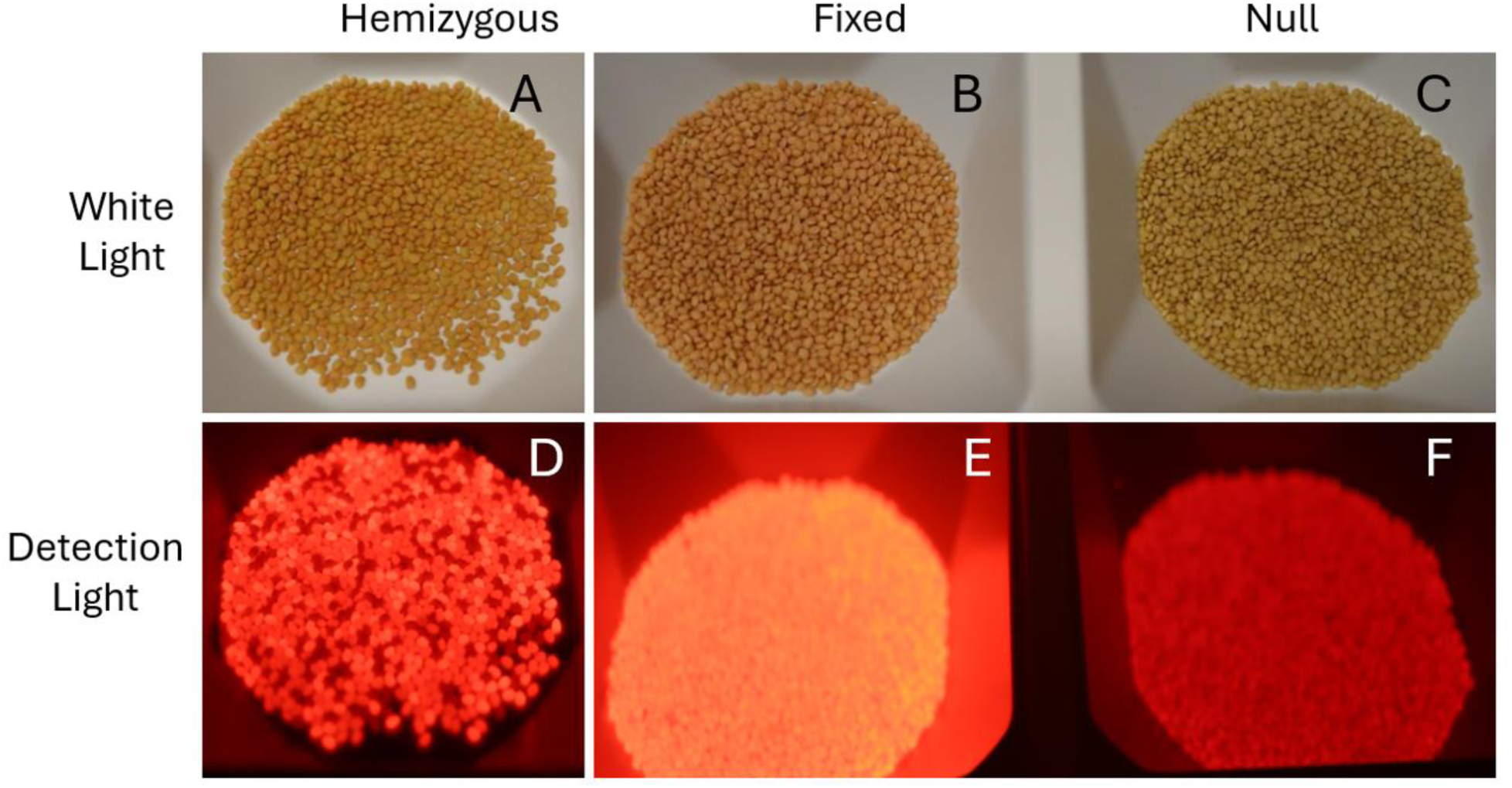
Illustration of DSRed2 expression differentiation among hemizygous, homozygous fixed, and homozygous null plant genotypes. Figures A and D demonstrate a segregating family in white light and under the detection light system, respectively. Figures B and E demonstrate a high intensity homozygous fixed family, and Figures C and F demonstrate a low intensity homozygous null family.

Selected seeds were planted, vernalized, and transplanted into 3.00 Kord Traditional Square pots. At the onset of flowering, these plants were arranged in a checkerboard configuration in a 3.00 Square Injection Pot 28 Pocket Kord Trays so that each plant neighbored plants of the opposite genotype. The canopy of each plant touched the canopy of all neighboring plants. Seed from each plant was harvested and threshed into single plant packets. The resulting seed was visualized under a detection light system (PSI SL3500-D-Green DSRed) to ascertain the presence of hemizygous seeds that would indicate outcrossing between the homozygous fixed and homozygous null plants.

#### 2.2.2 Cross-pollination method study

A second greenhouse experiment evaluated outcrossing rates under artificial conditions that mimic possible pathways for cross pollination in field situations. The pollen donor was CoverCress variety B3:WG (tt8/fae1), which is fixed for recessive mutations of transparent testa 8 (tt*8*) and fatty acid elongase 1 (*fae1*). The pollen receptor was CoverCress B3 wild type (WT, fixed dominant for TT8 and FAE1).

Table 1 summarizes the two control and three experimental treatments. Treatments 1 and 2 were negative and positive controls, respectively. Treatment 1 consisted of both B3:WG and B3 wild type plants produced at the same time but physically isolated in the greenhouse and allowed to naturally pollinate.

**Table 1.** Summary of control and experimental treatments in the greenhouse cross pollination study.

| Treatment | Protocol |
| --- | --- |
| 1. Negative Control | 8 WT (fixed FAE1 and TT8) and 8 WG (fixed fae1 and tt8) plants grown separately as negative control |
| 2. Positive Control, Forced Cross-Pollination | 4 WT female and 4 WG male plants were crossed as a positive forced pollination control |
| 3. Pollination Without Emasculation | The flowers of 4 WT female plants were touched by 4 WG male flowers manually in the morning in the flowering period to potentially mimic insect pollination |
| 4. Pollination With Emasculation | The emasculated flowers of 4 WT female plants were touched by 4 WG male flowers manually in the morning in the flowering period as a bee mimic treatment |
| 5. Intensive Flower Interaction | 14 WT and 14 WG plants were planted side by side. At peak pollination, they were covered with a plastic bag and shaken vigorously to promote pollen distribution to mimic potential wind pollination |

Treatment 2 used B3 wild type plants as receptor plants when pollen from B3:WG plants were used for forced cross-pollination. The cross pollination was performed by emasculating unopened flower buds on receptor plants at a timing before self-pollination was expected to occur (approximately 10 days before flower opening) and physically moving pollen from B3:WG plants to B3 wild type plant flowers.

Receptor plants in treatments 2 and 4 were emasculated at the same timing. Treatment 3 (pollination without emasculation) brought intact open receptor flowers in direct contact with actively pollinating donor flowers to create an enhanced natural pollination condition. Treatment 4 (pollination with emasculation) brought emasculated receptor flowers into direct contact with an actively pollinating donor flower, mimicking the potential for insect pollination (though more severe as a result of emasculation).

Treatment 5 (intensive flower interaction) placed receptor and donor plants in alternating cells of a tray in a checkerboard pattern and enclosed them in a plastic covering, then subjected them to enhanced pollen movement through agitation at peak flowering to mimic wind pollination. The plastic covering was removed after the treatment. Plants were grown in HC 3.00 Traditional Square Pot (Dimension: 7.62 × 7.62 × 6.35 cm) in 3.00 Square Injection Pot 28 Pocket Kord tray (Length / Width/Height: 52.71 cm × 30.48 cm × 5.72 cm) to maturity, and seeds from each plant were harvested separately into a 50-seed sample plus a remnant seed packet. qPCR identified the presence or absence of the fae1 allele in each sample.

### 2.3. Field Evaluation of Domesticated Pennycress Pollen Flow

A field-based pollen flow study in the 2023-2024 growing season using a Nelder Wheel design (Mark, 1983; Maity et. al. 2022) provided data on the potential for pennycress outcrossing under field conditions. The design included a 7.6 m diameter circle sown with the pollen donor variety (CoverCress B3:WG, fixed tt8, fae1) at a rate of 5.6 kg/ha (Figure 3). Eight uniformly spaced transects radiated from the central circle with 1 m single row pollen receptor plots spaced every 1.5 m along each transect. Each pollen receptor plot was sown with 30 seeds of CoverCress variety B3:LF (fixed tt8 and FAE1). Ten to twenty plants were established in each plot and grown to maturity. Presence of the fae1 allele in seed from pollen receptor plants is expected only if it has received pollen from a plant in the B3:WG central donor plot. The fae1 mutation was tracked with molecular markers. Frequency of the fae1 allele in pollen receptor plots indicated the distance and direction of pollen movement.

**Figure 3.**
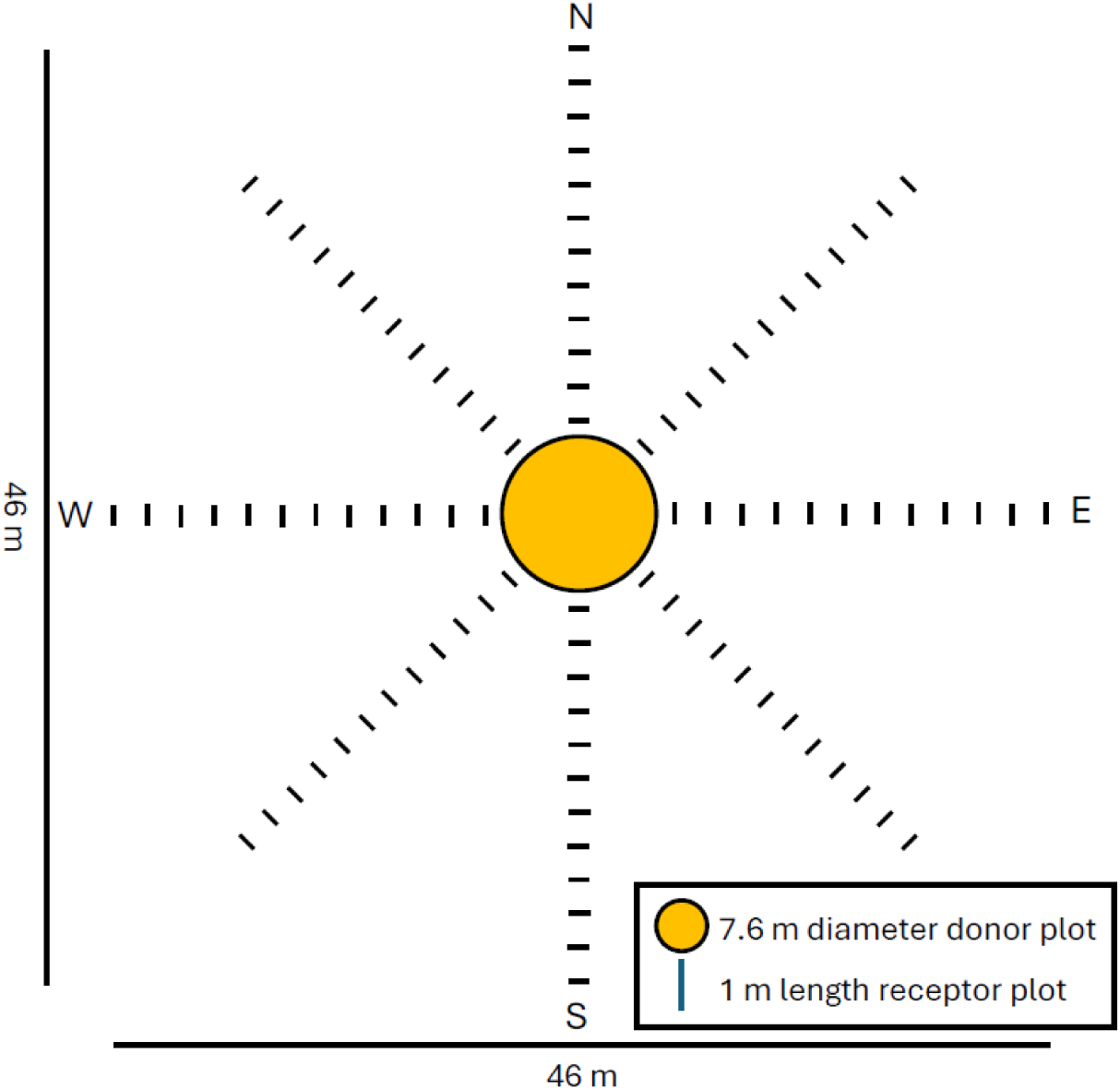
Nelder Wheel design of field pollen flow trial. Center donor plot comprised of B3:WG (fixed tt8 and fae) is 7.6 m diameter, surrounded by 1 m receptor plots of B3:LF (fixed tt8) spaced every 1.6m along lateral intersects as shown. Predominant winds flow from west to east.

The two trial locations were near Mascoutah, Illinois and at the Donald Danforth Plant Science Center Field Research Farm, near St. Charles, Missouri. Both sites have Mollisol soil types and are moderately well drained and well drained, respectively. The average annual precipitation for Mascoutah and St. Charles was 111.3 cm and 111.7 cm, respectively. Sites received two vertical tillage passes, split nitrogen applications in the fall and spring (56 kg/ha (50 lbs/acre) total), and a pre-emergence trifluralin herbicide application (3.2 L/ha).

Domesticated pennycress grown under field conditions typically flowers over a period of three to four weeks (first open flower to flower completion). Donor and receptor flowering reached 50% flowering (half of plants expressing at least one open flower) on March 28, 2024, and April 5, 2024, at Mascoutah and St. Charles, respectively. Temperature and precipitation data were collected from the National Oceanic and Atmospheric Administration (NOAA) weather station nearest to each research site. The average daily temperature during the flowering periods was 12°C and 13°C for the Mascoutah and St. Charles locations, respectively. Intermittent peak temperatures above 30°C, which are potentially detrimental to successful pollination, occurred at both locations toward the end of the flowering window. The 2023-2024 growing season was influenced by an “El Niño” weather pattern that began in May 2023 and lasted until May 2024. Above normal temperatures occurred in eight out of nine months of the growing season at both locations and average temperatures were 1.5°C and 1.8°C above long-term averages at Mascoutah and St. Charles, respectively. Seed set in the plots was typical for domesticated pennycress was normal despite the elevated temperatures. The overall temperature pattern would not have precluded cross-pollination throughout the season.

Weed pressure was present at both trial locations. Henbit (*Lamium amplexicaule*, L.), chickweed (*Stellaria media*, L.), chamomile (*Matricaria chamomilla*, L.), native field pennycress and shepherds’ purse (*Capsella bursa-pastoris*, L.) were the most common weeds. Weeds were mowed after CoverCress flowering was completed and pod grain filling was in progress. The presence and movement of pollinators within the trial was not evaluated quantitatively as part of the study, but study personnel observed butterflies and other pollinating insects at both locations.

Plot harvest was taken place on May 13, 2024 (Mascoutah) and May 21, 2024 (St. Charles) when plants reached full maturity and seed moisture declined to 8-12%. Harvest consisted of collecting pods from the main stem of five plants selected randomly (one plant from each end of the plot and three from the middle) across each of the 96 receptor plots at each of the two locations. Harvested pods from individual plants were threshed manually with gloved hands and seed was collected in individual packets. Donor plot seed samples were from the main stem of random plants from the middle of the donor plot and at the edge near the receptor plots. qPCR testing of seed aliquots from both donor and receptor plots confirmed the presence or absence of outcrossing.

### 2.4. Molecular Assay

A quantitative bulk seed assay was developed to detect seed from receptor plants that were cross-pollinated by the fae1 allele from the pollen donor. Self-pollinated receptor seed expresses the homozygous genotype FAE1, while outcrossed seed expresses the heterozygous FAE1/fae1 genotype. The assay consisted of extracting DNA from each 50 seed pool using a Quick-DNA Plant/Seed Miniprep kit made by ZYMO Research followed by an allele-specific assay for the fae1 and FAE1 alleles.

Sensitivity of the quantitative assay for fae1 was evaluated using eleven pools of 50 seeds with varying percentages of homozygous fae1 seed (0%, 2%, 4%, 6%, 8%, 50%, 92%, 94%, 96%, 98%, and 100%). DNA from the seed pools was extracted using the ZYMO Quick-DNA Plant/Seed Miniprep Kit and analyzed with fae1 primers on a Quant Studio 6 thermal cycler (Thermo Fisher Scientific). Figure 4 presents the results of this analysis, demonstrating that the assay successfully differentiated between seed pools with as little as a 2% difference (1 seed in 50) in the ratio of one fae1/fae1 to 49 FAE1/FAE1 seed mixture. Outcrossing in the field trial would produce a heterozygous fae1/FAE1 genotype rather than the homozygous genotype used in the calibration, requiring a 2x adjustment to the percentage seed predicted by the model. These results suggest the assay is capable of detecting outcrossing as low as 2% homozygous contaminant or 4% heterozygous seed in pools of 50 seed.

**Figure 4.**
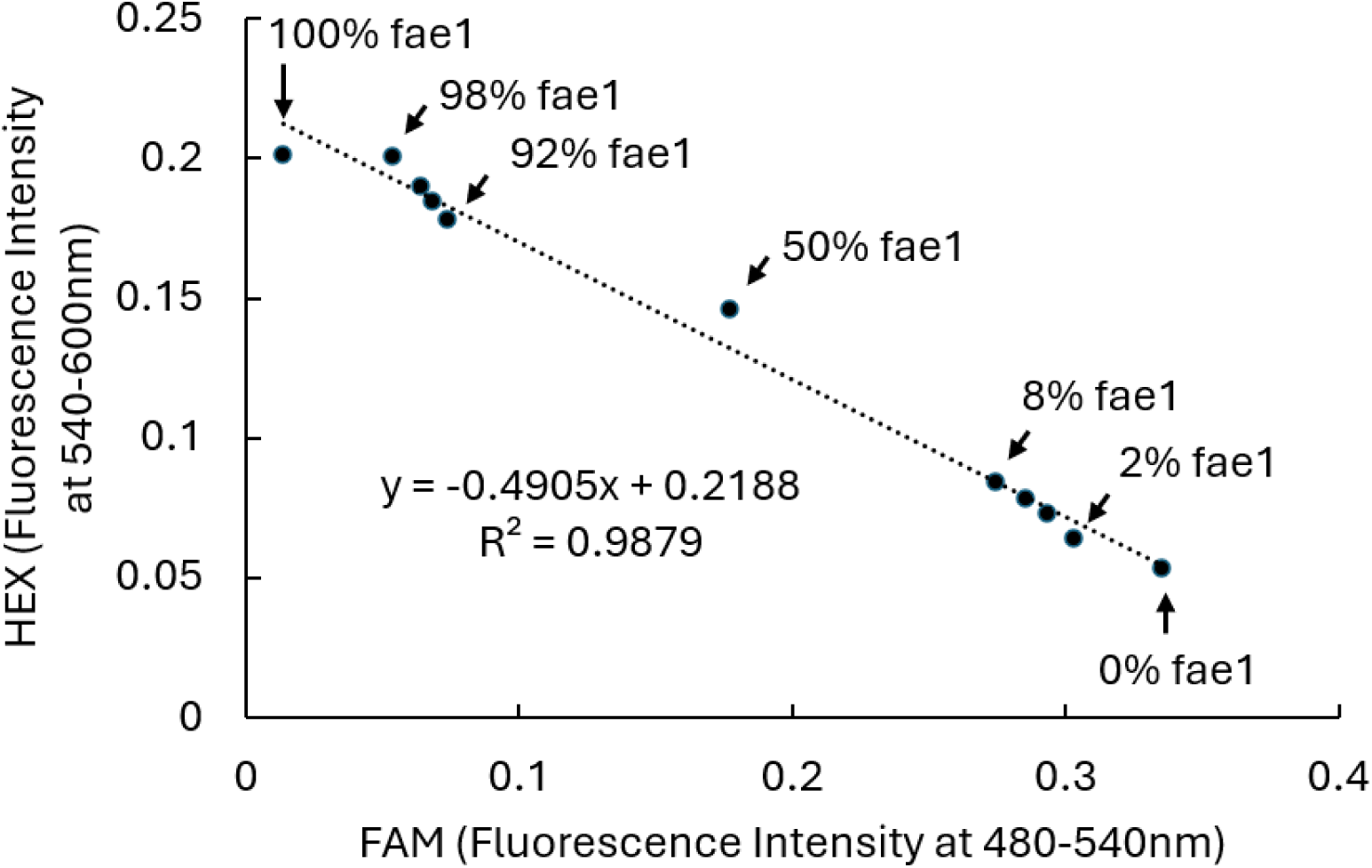
Reference data scatter plot demonstrates ability to detect single seed differences in homozygous fae1 contamination. Pools of 50 predominantly wild type homozygous FAE1 seeds were tested. A single homozygous fae1 seed contamination indicates a difference of 2%. This corresponds to 2 heterozygous fae1/FAE1 seed in a pool of 50 seeds (difference of 3.9%).

## 3. Results

### 3.1 Pollen viability kinetics

Domesticated pennycress pollen grains were spheroidal in shape and measured approximately 18 µm × 25 µm. As expected, viability decreased with increasing treatment exposure time, increasing temperature, and exposure to sunlight (Figure 5). Pollen longevity (T_1/2_ or number of hours to reach 50% of initial viability) was highest in the 4°C treatment, where it reached zero after 36 h exposure. The shortest T_1/2_ of 1.6 h was observed in samples placed outside and exposed to natural sunlight, wind, and variable spring temperatures. Under these conditions, pollen viability dropped to 4% after just 3 h exposure and reached zero after 8 h. Indoor temperature treatments demonstrated a longevity reduction with increasing temperatures with T_1/2_ of 12 h, 6 h, and 2.5 h for 4°C, 22°C, and 37°C, respectively. A pollen viability kinetics curve (Figure 5) characterized the interaction of temperature and time on declines in pollen viability. Note that moisture, wind, and light differences are confounded across treatments.

**Figure 5.**
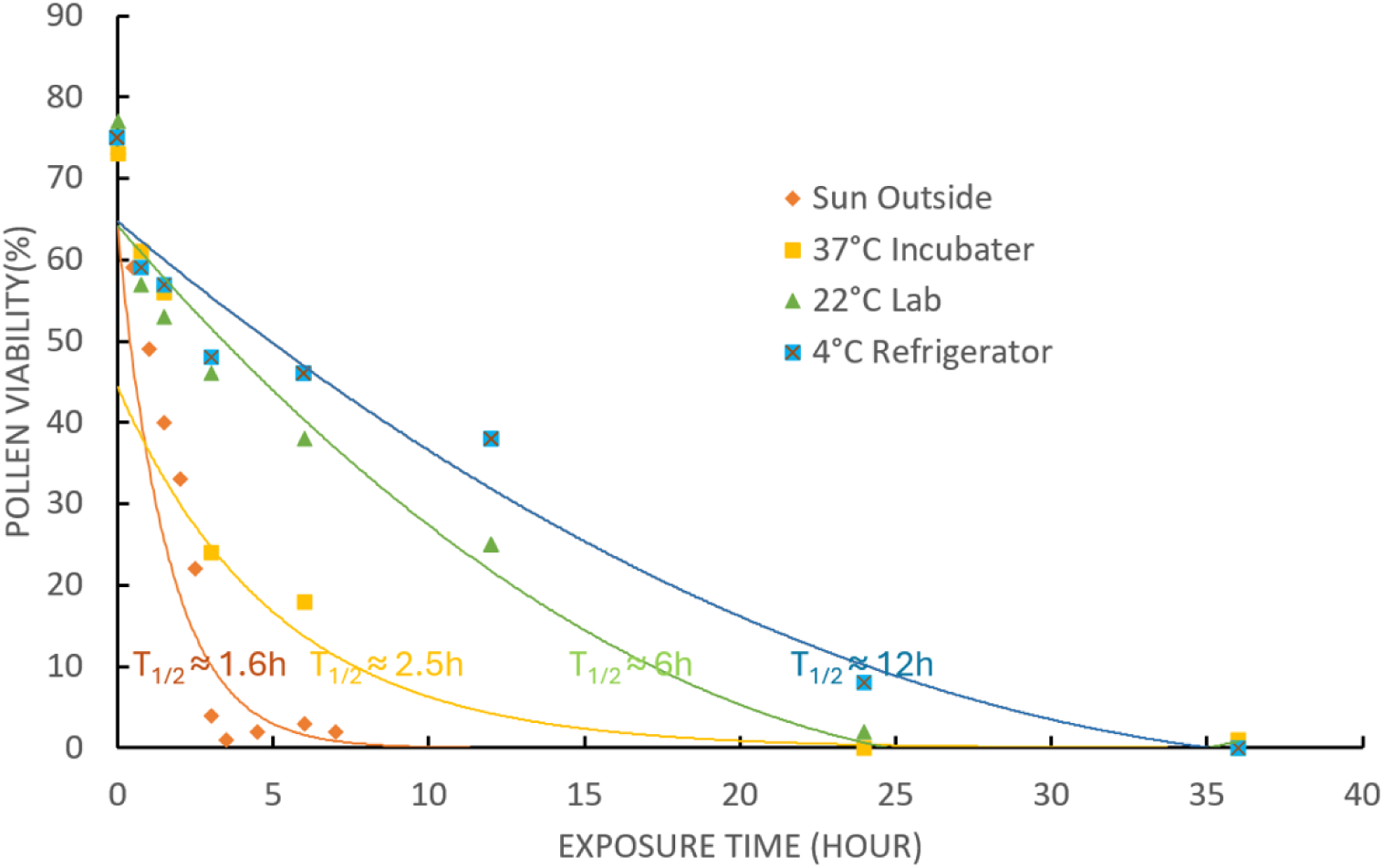
Pollen viability measured by fluorescein diacetate (FDA) fluorescence of domesticated pennycress after various exposure times under different temperatures and environmental conditions. Viability declined over time under all tested conditions (Orange, outside under sun; Yellow, 37 °C incubator; Green, 22 °C lab; Blue, 4 °C refrigerator) and fell the most rapidly when exposed to direct sunlight. T1/2 is the time at which 50% of pollen was found to be inviable. Increasing temperatures were associated with lower T1/2 values, demonstrating that length of pollen viability decreases with increasing temperature.

### 3.2. Greenhouse Outcrossing Trials

#### 3.2.1. Close proximity cross-pollination

All harvested seed exhibited the DsRed expression level of the parental seed, indicating absence of cross-pollination among expression groups (Table 2). None of the harvested seeds exhibited the intermediate DsRed expression phenotype that would indicate outcrossing between DsRed expressing and DsRed non-expressing plants. No DSRed2 expression was detected in any of the seed lots of the close-proximity cross-pollination study harvested from null T2 plants (Table 2). This result indicates that no outcrossing occurred among plants despite their floral organs remaining in close proximity in greenhouse conditions.

**Table 2:**
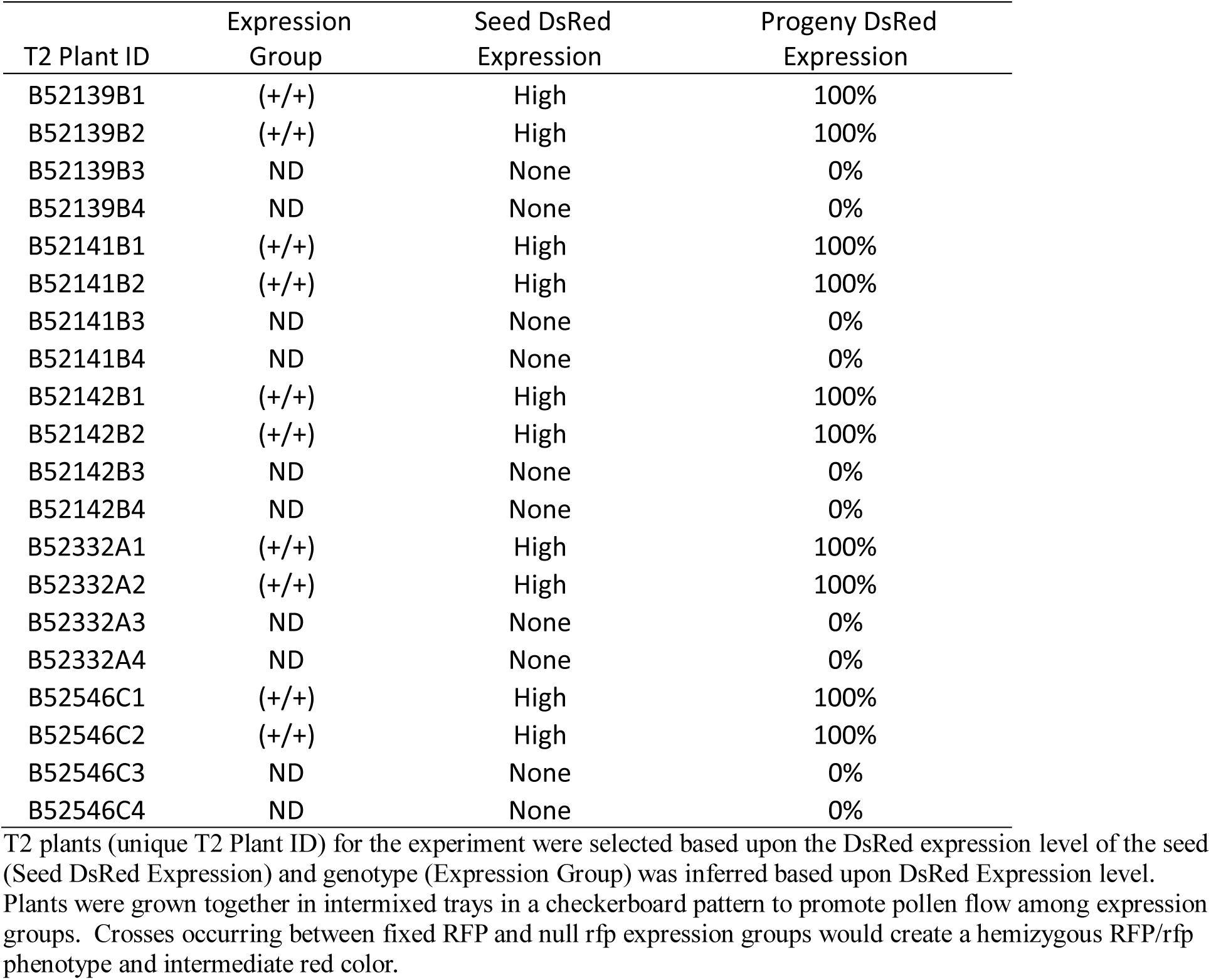
Results of the close-proximity cross-pollination study. T2 plants (unique T2 Plant ID) for the experiment were selected based upon the DsRed expression level of the seed (Seed DsRed Expression) and genotype (Expression Group) was inferred based upon DsRed Expression level. Plants were grown together in intermixed trays in a checkerboard pattern to promote pollen flow among expression groups. Crosses occurring between fixed RFP and null rfp expression groups would create a hemizygous RFP/rfp phenotype and intermediate red color.

#### 3.2.2 Cross-pollination method study

The second greenhouse trial included two controls and three experimental treatments with varying levels of susceptibility to cross-pollination. As expected, the negative control produced no detectable outcrossing, and the positive control produced 100% outcrossing. The Pollination with Emasculation treatment (Treatment 4) was the most extreme and produced the highest rate of cross-pollination among the three experimental treatments (36%; 40 of 110 harvested seeds) (Table 3). In the Intensive Flower Interaction (Treatment 5), one of 24 seed pools (50 seeds per pool) exhibited detectable outcrossing. The outcrossing rate of the Intensive Flower Interaction study was therefore approximately 0.2% (2 × 1 outcross / (24 pools × 50 seed/pool) = 0.2%). No outcrossed seed was detected in the Pollination Without Emasculation treatment (Treatment 3) in which a single pool of 45 seeds were tested. These results suggest the rate of outcrossing in pennycress is low, particularly given the low rate observed in the Intensive Flower Interaction treatment.

**Table 3.** Outcrossing rates based on molecular assay using pools of 50 seed in greenhouse-controlled conditions. Negative control consisted of plants grown in isolation. The positive control involved forced pollination where WT (Female) was crossed to WG (male). Pollination Without Emasculation was similar to the negative control and no outcrossing was observed. Pollination With Emasculation was intended to mimic the effects of pollinating insects and showed outcrossing in 36% of the 110 harvested seeds. The Intensive Flower Interaction was intended to create outcrossing pressure similar to what might be caused by wind. Seed from all treatments were subjected to qPCR assays to determine the cross-pollination rate.

| Treatment | # Seeds Analyzed | Outcrossing Rate (%) |
| --- | --- | --- |
| 1. Negative Control | 400 (8 pools) | Not detected |
| 2. Positive Control, Forced Cross-Pollination | 15 | 100 |
| 3. Pollination Without Emasculation | 45 | Not detected |
| 4. Pollination With Emasculation | 110 | 36 |
| 5. Intensive Flower Interaction | 1200 (24 pools) | <2 |

### 3.3 Field evaluation of domesticated pennycress pollen flow

Nine sub-samples (450 seeds) were tested from the WG pollen donor plot in each location. These samples all indicated a 100% fae1/fae1 genotype, confirming the fae1 genotype was fixed in the plot and indicating that no detectable outcrossing with pollen from receptor plots occurred. Molecular testing to examine FAE1/fae1 frequency in the four LF (receptor) plots closest to the donor plot (1.5 m, 3 m, 4.5 m, 6 m from the donor plot) in all eight directional transects produced no evidence of heterozygous, outcrossed FAE1/fae1 seed, indicating outcrossing was undetectable. Two 50-seed sub-samples (100 seeds total) from each of the 32 LF (receptor) plots at both field locations were assayed for a total of 128 assays.

Despite not observing evidence of pollen flow from the nearest receptor plots, plots 9 m and 18 m, from the pollen donor, in all eight directional transects for both field locations were tested. All 64 replicates tested again showed no evidence of outcrossing. Collectively, pollen flow data from both experimental fields suggest a very low rate of outcrossing, below the detection limit of the assay.

## 4. Discussion

Pennycress plants are diploid and reproduce predominantly through self-fertilization (Best and McIntyre, 1975; Warwick et al., 2002; Dorn et al., 2015), though it has been suggested that wind (Groeneveld and Klein, 2014) or insect pollination (Al-Shehbaz, 1986) may also play a role. This study presents the first evaluation of pollen flow in field environments using a trackable indicator result and supports these results with additional description of pollen viability and movement in controlled environments. All results indicate that domesticated pennycress is predominantly self-pollinating with very low rates of outcrossing.

The floral morphology of pennycress, consisting of six stamens with four possessing anthers at the same height or slightly higher than the stigma, makes pennycress readily autogamous as the stamens curl inward as flowers mature (Warwick et al., 2002; Best and McIntyre, 1975). Targeted cross-pollination in CoverCress Inc. research programs must account for the fact that most pennycress flowers are already self-fertilized before opening. To successfully cross pennycress plants, breeders emasculate floral buds about ten days before flower opening to prevent self-pollination (CoverCress Inc., unpublished data).

This observation strongly supports self-pollination as the predominant means of reproduction in pennycress.

Laboratory experiments exposing domesticated pennycress pollen to elevated temperatures or sunlight confirmed that both stressors can reduce viability, supporting a conclusion that environmental exposures limit the ability of pennycress pollen to travel and successfully fertilize other pennycress plants. Similar to the present studies, Parker and Phippen (2012) reported an 84% reduction in pennycress pollen viability at 30°C and a 100% reduction at 32°C. Consistent with these observations, Zhang et al. (2020) reported that pollen from *Camelina sativa*, L. (camelina), a species related to pennycress, has a half-life life (T_1/2_) of 0.4 hours as evaluated through pollen grain germination and that no camelina pollen remained viable after five hours under ambient conditions (Zhang et al., 2020). Similarly, Gray et al. (2024) reported that UV-B light reduced pollen viability after anthesis in *Brassica rapa*, L., a fellow member of the Brassicaceae with pennycress. Likewise, Cun et al. (2024) concluded that UV-B light reduces pollen viability in a meta-analysis of 77 species in 30 families. The authors reported pollen from 90.9% and 84.2% of studied species exhibited reduced pollen germination or pollen tube growth, respectively in response to UV-B light.

Data from the present study demonstrates a T_1/2_ of 1.6 hours for domesticated pennycress when pollen was placed outside and exposed to natural sunlight with wind. While this seems considerably longer than the T_1/2_ of camelina (0.4 hours), the FDA analysis has been shown to overestimate pollen longevity compared to in vitro germination tests in the camelina evaluation (Impe et al., 2020), suggesting that the T_1/2_ for domesticated pennycress pollen viability via germination may be shorter than the observed 1.6 hours and closer to camelina. The warm temperatures and potential exposure to sun required for insect activity (Jaboor et al., 2022) likely reduce the length of pollen viability and lower the frequency of successful cross-pollination, even when pollinating insects are present.

Greenhouse experiments confirmed that the potential for outcrossing in pennycress is low. In the close-proximity cross-pollination study, domesticated pennycress plants growing in contact with domesticated pennycress plants expressing the DsRed selectable marker showed no detectable evidence of outcrossing. In the cross-pollination method study, only the positive control and the Pollination with Emasculation treatment, where emasculated plants came into direct contact with anthers from pollen donor plants, produced detectable cross pollination.

Field experiments provided further confirmation that pennycress is unlikely to undergo significant outcrossing in real-world conditions. Nelder wheel plots grown at two locations in geographies where domesticated pennycress is grown produced in no detectable cross pollination at distances as close as 1.5 m between pollen donor and pollen receptor plants. These results support the hypothesis that pennycress cross-pollination under field conditions is rare even over short distances and that pennycress is predominantly self-pollinated.

The molecular-based assay to detect outcrossing and seed lot contamination uses a quantitative PCR approach to detect the amount of *fae1* mutant genotype present in a 50 seed pool. The assay can detect a single homozygous seed or two heterozygous seeds out of a pool of 50 seeds. Most existing methods for similar purposes have detection limits that prevent resolution below about 1% (Nageswara-Rao et al. 2013; Xu et al. 2021). During development, it was clear that this level of resolution was not achievable as a result of limitations in seed grinding and PCR sensitivity for the marker technology. Despite the higher 4% detection limit for outcrosses in a single pool, greater accuracy can be achieved by evaluating multiple pools. Relying on multiple lines of evidence from laboratory, greenhouse, and field experiments, however, mitigates the potential impact of this limitation.

## Acknowledgements

The authors acknowledge the support of Anastasia Shamin in the execution of molecular assays, Tyler Mueth for support on field trial layout, seeding, and management, Paige Perry for data generation on the pollen viability assay, Dr. Michelle Starke for reviewing and commenting the manuscript and Hjelle Consulting Group, LLC for editorial assistance.

## Author Contributions

T Lavaire. Designed and performed the field research trial.

D McLaughlin. Contributed to development of molecular analysis tools and performed the research.

S Liu. Designed and performed pollen viability kinetics and cross-pollination methods experiments; analyzed data, wrote significant portions of the paper and reviewed the paper.

R Kennedy. Designed and performed a close-proximity cross-pollination study and wrote the description.

T Sauer. Performed cross-pollination experiments and contributed to the molecular validation of the experiments.

R Chopra. Designed the molecular assays and contributed to trial design and execution, wrote and reviewed the paper.

K Cook. Conceived the idea and led the execution of this work, contributed to field trial design and execution; interpreted analysis, and wrote the paper.

Statement of conflict: All authors are employees of CoverCress, Inc. who are domesticating field pennycress as a new winter intermediate oilseed crop and have filed patents in the crop. These patents are unrelated to this manuscript.

